# A Peptide of the Amino-Terminus of GRK2 Induces Hypertrophy and Yet Elicits Cardioprotection after Pressure Overload

**DOI:** 10.1101/2020.06.18.159194

**Authors:** Sarah M. Schumacher, Kamila M. Bledzka, Jessica Grondolsky, Rajika Roy, Erhe Gao, J. Kurt Chuprun, Walter J. Koch

## Abstract

G protein-coupled receptor (GPCR) kinase 2 (GRK2) expression and activity are elevated early on in response to several forms of cardiovascular stress and are a hallmark of heart failure. Interestingly, though, in addition to its well-characterized role in regulating GPCRs, mounting evidence suggests a GRK2 “interactome” that underlies a great diversity in its functional roles. Several such GRK2 interacting partners are important for adaptive and maladaptive myocyte growth; therefore, an understanding of domain-specific interactions with signaling and regulatory molecules could lead to novel targets for heart failure therapy. While elevated cardiac levels and activity of GRK2 contribute to adverse heart remodeling and contractile dysfunction, inhibition of GRK2 via overexpression of a carboxyl-terminal peptide, βARKct, or its amino-terminal domain Regulator of G protein Signaling (RGS) homology domain (βARKrgs) can enhance cardiac function and can prevent heart failure development via G_βγ_ or Gαq sequestration, respectively. Previously, our lab investigated cardiac-specific transgenic expression of a fragment of this RGS domain (βARKnt) (residues 50-145). In contrast to βARKrgs this fragment did not alter acute hypertrophy after pressure overload or demonstrate RGS activity *in vivo* against Gq-mediated signaling. Herein, we subjected these transgenic mice to pressure overload and found that unlike their littermate controls or previous GRK2 fragments, they exhibited an increased left ventricular wall thickness and mass prior to cardiac stress that underwent proportional hypertrophic growth to controls after acute pressure overload. Importantly, despite this enlarged heart, βARKnt mice did not undergo the expected transition to heart failure observed in controls. Further, βARKnt expression limited adverse left ventricular remodeling and increased cell survival signaling. These data support the idea that the βARKnt peptide embodies a distinct functional interaction and novel means of cardioprotection during pressure-overload induced heart failure.

## 1. Introduction

G protein-coupled receptor (CPCR) kinase 2 (GRK2) is the subject of a growing body of research not only into cardiac function, but diverse chronic and systemic diseases. In addition to its well characterized role in the regulation of GPCRs, this ongoing research is identifying new and diverse functional roles for GRK2 in cellular regulation. While many of these GRK2 functions have been described, the mechanisms by which they occur *in vivo* are poorly understood, often occurring through kinase-independent scaffolding and protein interaction mechanisms. This may be due to our limited understanding of the distinct functional domains of GRK2, the central catalytic domain and amino and carboxyl-terminal regulatory domains, and the distinct protein interactions they encompass. One such domain is the amino terminal <u>R</u>egulator of <u>G</u> protein signaling or RGS homology domain (amino acids 51-173), that we have previously shown can selectively interact with Gαq in vivo [1]. This interaction sequestered Gαq, preventing downstream signaling through Gq-coupled GPCRs and thus reducing hypertrophy and preventing heart failure [1].

It is clear from the literature that GRK2 has a robust interactome and its activity can affect several substrates including non-GPCR targets [2, 3]. Although we have previously shown that the RGS domain of GRK2 can interact with Gαq [1] and the carboxyl terminus (βARKct) can interact with G_βγ_ and also Heat shock protein (Hsp) 90 [4–6]; it isn’t clear if other domains and other peptide regions target other interacting partners. Previously, our lab investigated transgenic mice with cardiac-specific expression of a fragment of GRK2’s RGS domain (residues 50-145), which we have termed βARKnt. This smaller amino-terminal peptide of GRK2 induced baseline hypertrophy and elevated β-adrenergic receptor (βAR) density without altering downstream signaling [7]. Since the βARKnt does not incorporate the entire RGS domain, it expectedly did not interfere with myocardial Gαq signaling [7, 8]. In order to determine the mechanism of action of this peptide and investigate whether the regulation of βAR density could alter cardiac disease progression, we generated new Tg mice with cardiac expression of the βARKnt peptide that contains a carboxyl-terminal flag tag for easier identification separate from any endogenous GRK2. In this study, we have stressed these TgβARKnt mice with left ventricular (LV) pressure overload via transverse aortic constriction (TAC) to determine whether there were specific effects of this peptide distinct from other GRK2 domains. Indeed, our data indicate specific effects on cardiac hypertrophy and dysfunction after TAC suggesting novel interacting partners of this peptide region of GRK2.

## 2. Materials and methods

### 2.1 Study Design & Experimental Animals

This study will focus on the use of 8-10 week old male laboratory mice for all proposed experiments. Females were not excluded from this study, rather their results were quite divergent and due to the nature of divergence will be the subject of a separate manuscript. The lines used in this proposal are all driven by the cardiac-restricted promoter α-myosin heavy chain (αMHC) and are on a C57BL/6J background. Cardiac-specific transgenic (Tg) mice expressing a flag-tagged βARKnt peptide were generated as described previously [7]. Briefly, the cDNA that encodes bovine GRK2 residues 50-145 was cloned into a vector driven by the αMHC promoter with a carboxyl-terminal flag-tag. All transgenic mouse studies include equal numbers of non-transgenic littermate control (NLC) mice for all planned experiments. Mice with αMHC-targeted expression of the βARKct peptide (CT 194 residues) containing the G_βγ_-binding domain were utilized as a control for GRK2 inhibition, while mice with αMHC-targeted expression of GRK2, causing a 2-3 fold increase in protein levels similar to the increase observed during human heart failure, were utilized as a control for enhanced GRK2 activity.

Statistical powering was performed using the G*Power 3.1.9.2 software from the University of Dusseldorf for power analysis and estimation of sample size. Based on these calculations our target was a minimum of 9 animals (powered for indices of trans-aortic constriction) per group to attain statistical significance, with a preference for 10-15. In general, separate cohorts of mice were used for, histology, biochemistry, membrane receptor density studies, and proteomic analysis. Each experiment was performed a minimum of 3 times. The number and composition of the replicates were determined based upon availability of transgenic and non-transgenic animals, the surgeon, and echocardiography and hemodynamic machinery. All animals and resulting samples were monitored by mouse number only until data quantification was complete and then decoded by gene expression and surgical group for statistical analysis, meaning all data analysis was blinded. All results were substantiated by repetition. Data were only excluded if their validity was undermined by the condition of the animal or cells prior or during the experiment, such as loss of the specimen. All animal procedures were carried out according to National Institutes of Health Guidelines on the Use of Laboratory Animals and approved by the Animal Care and Use Committees of Temple University and the Lerner Research Institute (LRI).

### 2.2. In vivo Hemodynamics

*In vivo* cardiac hemodynamic function was measured at baseline and after administration of the βAR agonist isoproterenol (0.1, 0.5, 1, 5, and 10 ng) as described [9].

### 2.3. In vivo TAC Model & Echocardiography

Transverse aortic constriction (TAC) was performed as described previously [1, 10]. Echocardiography was performed using the Vevo 2100 imaging system from VisualSonics as described [11]. Briefly, two-dimensional echocardiographic views of the mid-ventricular short axis were obtained at the level of the papillary muscle tips below the mitral valve. M-mode measurements were determined at the plane bisecting the papillary muscles according to the American Society of Echocardiography leading edge method. To measure global cardiac function, echocardiography was performed at 8-10 weeks of age prior to TAC and at 2 and 4, or 2, 4, 6, 8, 10, 12 and 14 weeks post-TAC. Pressure gradients were determined 1 week later by pulsed-wave Doppler echocardiography of the transverse aorta and mice with gradients over 50 mmHg were used. Subgroups of hearts were harvested at 4 and 14 weeks after TAC for analyses of structure, fibrosis, and biochemistry.

### 2.4. RNA Isolation and Semi-quantitative PCR

RNA isolation and analysis was performed as previously described [1]. Total RNA isolation was performed using TRIzol reagent (Life Technologies) and a VCX 130PB ultrasonic processor from Sonics & Materials Inc (Newtown, CT) for homogenization. After RNA isolation, cDNA was synthesized from 1mg of total RNA using the iScript cDNA Synthesis Kit from BioRad Laboratories (Hercules, CA). Semi-quantitative PCR was carried out on cDNA using SYBR Green (Bio-Rad) and 150nM of gene-specific oligonucleotides for *18S, ANF, BNP, βMHC, MMP2, Col III, TGFβ* on a CFX96 Real Time System with BioRad CFX Manager 2.1 software (Bio-Rad). Quantitation was established by comparing 18s mRNA, which was similar between groups, for normalization and compared using the ΔΔCt method.

### 2.5. Histological Sectioning and Staining

Subgroups of hearts were harvested at 4 and 14 weeks post-TAC for analyses of structure, fibrosis, and biochemistry. Trichrome staining was performed as previously described [9]. Briefly, mice were euthanized and following cardioperfusion, hearts were fixed for 1-3 days in 4% paraformaldehyde at 4°C. Hearts were dehydrated and paraffinized using a Microm STP 120 from ThermoFisher Scientific, embedded in paraffin using a HistoStar apparatus (ThermoFisher), and sectioned (4-6 micron) using a Microm HM 325 (ThermoFisher). Tissue sections were then stained with Weigert’s iron hematoxylin and Masson Trichrome (Sigma-Aldrich) according to the manufacturer’s instructions. Interstitial fibrosis was quantified by color threshold measures using ImageJ. Alternatively, tissue sections were de-paraffinized and re-hydrated according to the Trichrome staining protocol, but following the wash with de-ionized water heart sections were washed 3 times 5 minutes with 1x PBS followed by staining with Alexa Fluor 488-conjugated wheat germ agglutinin (WGA, 1:10 in 1xPBS) (Invitrogen) for 1 hour at room temperature in a humidified chamber in the dark. Sections were again washed 3 times 5 minutes with 1x PBS followed by mounting with coverslips using Fluoromount-G mounting media containing DAPI nuclear stain (Southern Biotech).

### 2.6. Immunoprecipitation and Immunoblotting

Immunoprecipitation (IP) was performed as described previously [1]. Briefly, cardiac lysates were centrifuged at 13,000xg at 4° C for 30 min, protein concentration was measured using a BCA assay (Pierce), and 1mg of protein was used in the IP reaction. GRK2 and βARKnt-flag IPs were conducted using monoclonal anti-GRK2 agarose conjugate (Santa Cruz) and anti-Flag M2 affinity gel (Sigma-Aldrich), respectively. Control IPs were conducted using normal mouse IgG-agarose conjugates (Santa Cruz). Western blotting was performed as described previously [1]. Following SDS-PAGE and transfer to nitrocellulose membranes, primary antibody incubations were performed overnight at 4° C. Primary antibodies used were as follows: rabbit polyclonal or mouse monoclonal GRK2 (Santa Cruz), monoclonal GAPDH (Santa Cruz), monoclonal and polyclonal Flag (Sigma-Aldrich), and goat polyclonal Gαq/11 (Santa Cruz). Visualization of Western blot signals was performed using secondary antibodies coupled to Alexa Fluor 680 or 800 (Invitrogen Molecular Probes) or IRDye 800 (LI-COR Biosciences) and imaged using the Odyssey CLx infrared imager (LI-COR Biosciences) or ChemiDoc imaging system (Bio-Rad). Image Studio version 5.2 imaging software was used to process all images.

### 2.7. Inositol 1,4,5-trisphosphate Quantification

Inositol 1,4,5-trisphosphate (IP3) concentration was quantified as done previously [1] using a mouse IP3 ELISA kit according to the manufacturer’s instructions (MyBioSource).

### 2.8. Membrane Preparation and Radioligand Binding Assay for βARs

Myocardial sarcolemmal membranes were prepared by homogenization of whole heart left ventricular tissue as described [12]. Total βAR density was determined by incubation of 25 mg of cardiac sarcolemmal membranes with a saturating concentration (80 pmol/L) of [^125^I]cyanopindolol (CYP) and 20 mmol/L alprenolol to define nonspecific binding. Typical nonspecific binding is ~40% of the total. Competition-binding isotherms in sarcolemmal membranes were done in triplicate in 250 mL of binding buffer (50 mmol/L HEPES [pH 7.3], 5 mmol/L MgCl2, and 0.1 mmol/L ascorbic acid). Assays were conducted at 37°C for 60 minutes and then filtered over GF/C glass fiber filters (Whatman) that were washed and counted in a gamma counter and Kd and the maximal number of binding sites (Bmax) for ^125^I-CYP were determined by Scatchard analysis of saturation binding isotherms with GraphPad Prism as described [12].

### 2.9. Statistical Analysis

All values in the text and figures are presented as mean ± SEM of independent experiments for given n-sizes. Statistical significance was determined by one-way or two-way ANOVA with Tukey post-hoc test, Student’s *t* test, or nonparametric one-way ANOVA with Dunn’s post-hoc test as appropriate. For all statistical tests, a p value of < 0.05 was considered significant.

## 3. Results

### 3.1. βARKnt Transgenic Mice Exhibit Baseline Hypertrophy and Normal Adaptive Responses to TAC

To generate the TgβARKnt line, the cDNA that encodes bovine GRK2 residues 50-145 was cloned into a vector driven by the αMHC promoter with a carboxyl-terminal flag-tag. Founder lines were established and no gross phenotypic changes were observed in transgenic mice compared to non-transgenic littermate control (NLC) mice. Transgene expression was confirmed by Western blotting of cardiac lysates using a mouse monoclonal flag antibody to detect the ~15 kDa band (Fig. 1A). During initial characterization of the TgβARKnt line, cardiovascular function and adrenergic responsiveness were analyzed using terminal hemodynamics at baseline and upon challenge with increasing doses of the βAR agonist isoproterenol in transgenic and NLC mice. No difference was observed in mean systemic pressure (Fig. S1A), heart rate responses to isoproterenol (Fig. S1B), or cardiac contractility and relaxation (dP/dt maximum and minimum) at baseline and in response to isoproterenol (Fig. 1B, Fig. S1C), demonstrating that cardiac function and βAR responsiveness were not altered in our transgenic animals. This is consistent with previous data showing the βARKnt basally increases βAR membrane, but due to compensatory GRK2 levels, signaling was not enhanced [1].

**Figure 1:**
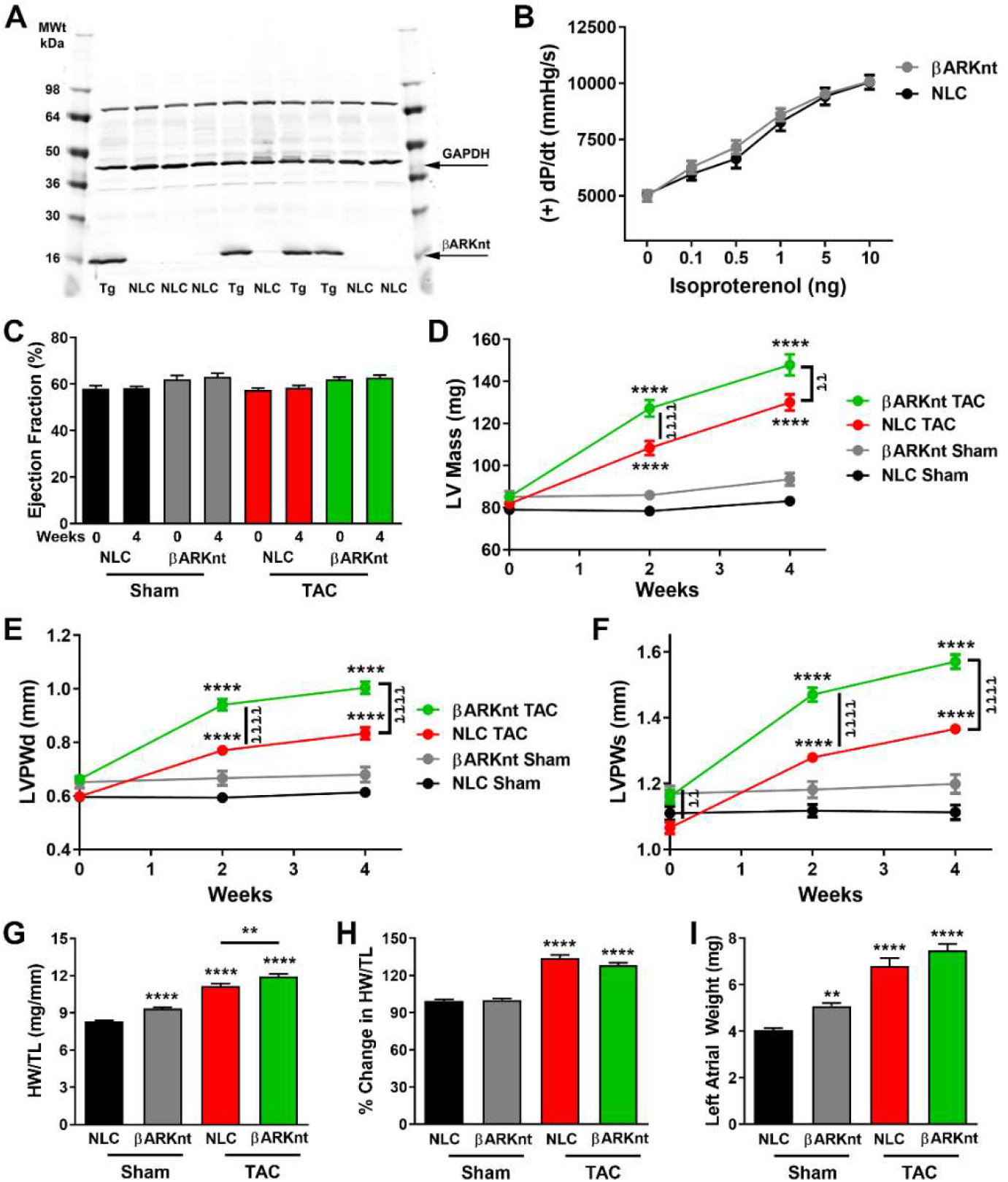
Cardiac-specific βARKnt expression elicits baseline hypertrophy and additive pressure overload-induced hypertrophy. (A) Western blot of TgβARKnt and non-transgenic littermate control (NLC) ventricular lysates probed for flag-tagged βARKnt and the loading control GAPDH. (B) Quantitation of LV +dP/dt average maximum from hemodynamics recordings of 10-12 week old TgβARKnt and NLC mice at baseline and with increasing doses of isoproterenol (0.1-10 ng). n = 12 mice per group. (C) % left ventricular (LV) ejection fraction from TgβARKnt and NLC Sham and post-TAC animals at baseline and 4 weeks post-surgery. Serial measures of noted experimental groups for (D) LV mass, (E) LV posterior wall thickness during diastole (LVPWd), and (F) systole (LVPWs). ****, p < 0.0001 by two-way ANOVA with repeated measures and Tukey post-hoc test relative to corresponding NLC Sham. ^ττ^, p ≤ 0.002; ^ττττ^, p < 0.0001 by twoway ANOVA with repeated measures and Tukey post-hoc test relative to corresponding NLC TAC. n = 9-17 mice per group. Measures of (G) heart weight normalized to tibia length (HW/TL), (H) % change in HW/TL, and (I) left atrial weight in these animals. **, p = 0.0083; ****, p < 0.0001 by one-way ANOVA with Tukey post-hoc test relative to NLC Sham. ^ττ^, p = 0.0094 by one-way ANOVA with Tukey post-hoc test relative to corresponding NLC TAC. n = 37-62 mice per group.

To determine whether expression of this peptide of GRK2 would alter cardiac remodeling in an animal model of pressure overload, TgβARKnt and NLC mice underwent TAC or sham surgery. Echocardiography was performed at baseline, 2, and 4 weeks to monitor cardiac function and dimensions. Sufficient LV pressure gradients after TAC were confirmed by pulsed-wave Doppler of the aortic arch 1 week post-surgery, with no differences between NLC and Tg groups (Fig. S1D). Cardiac function was assessed by measures of LV ejection fraction and fractional shortening (Fig. 1C, Table 1) that were not altered in either group over the 4 week time course. Interestingly, the increase in the LV mass after TAC, as well as LV posterior wall thickness during diastole and systole, were more significant in the TgβARKnt animals compared to NLCs (Fig. 1D-F). Accordingly, heart weight was greater in the βARKnt Sham and TAC mice 4 weeks after surgery (Fig. 1G); however, when normalized to their respective Sham animals, the % increase in HW after TAC was the same (Fig. 1H), and the increase in left atrial weight was equivalent (Fig. 1I), with no difference in tibia length (Fig. S1E). Together, these data demonstrate proportional hypertrophic growth in the βARKnt transgenic and NLC mice in response to cardiac stress.

**Table 1.**
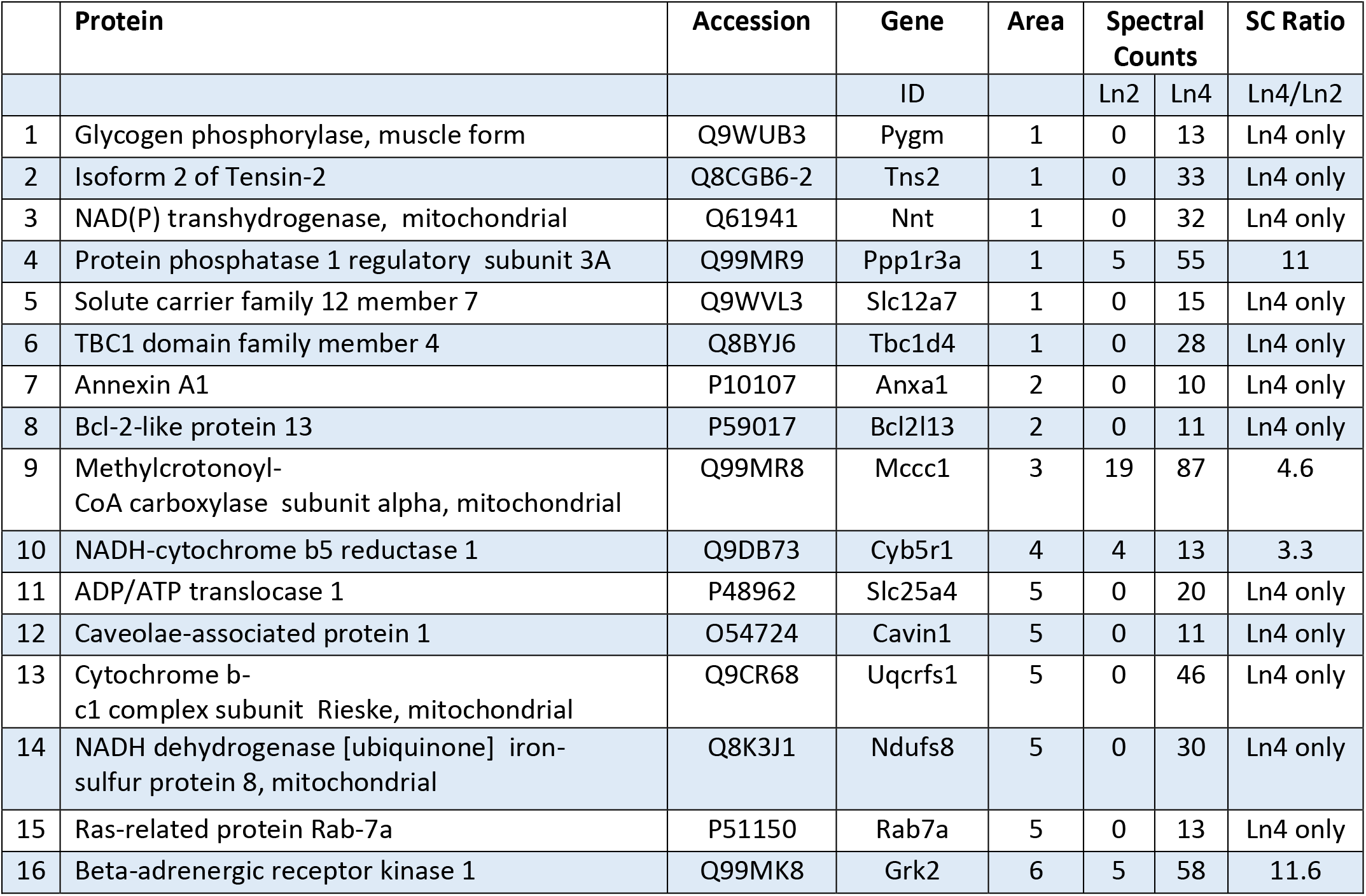
Summary for βARKnt-Flag IP

### 3.2. βARKnt Expression Protects Against Decompensated Hypertrophy and Heart Failure

To investigate whether the larger heart at baseline and after TAC induced by βARKnt expression would hasten the transition to heart failure, we followed a cohort of mice out to 14 weeks post-TAC. Cardiac structure and function were monitored by serial echocardiography every 2 weeks, and systolic pressure gradients were no different between the TAC groups 1 week postsurgery (Fig. S2A). As expected, NLC mice exhibit a robust transition to heart failure from 10-14 weeks, with a greater than 20% drop in ejection fraction by 14 weeks post-TAC (Fig. 2A) [1]. In contrast to this noticeable deterioration in function, ejection fraction was stable in the TgβARKnt mice throughout the entire time course similar to the Sham controls. While LV mass continually expanded in NLC mice, this measure was relatively flat in TgβARKnt mice from 4-14 weeks post-TAC, allowing the controls to achieve the same mass (Fig. 2B), with a similar trend observed in LV posterior wall thickness during diastole (Fig. S2B). Further, the progressive reduction in LV posterior wall thickness and expansion of the internal diameter during systole observed in the non-transgenic mice beginning at 8 weeks post-TAC, indexes of decompensation to heart failure, were abolished in the TgβARKnt mice (Fig. 2C, D). Consistent with these results, heart weight and left atrial weight normalized to tibia length were equivalent in the non-transgenic and βARKnt mice 14 weeks after TAC (Fig. 2E, F, G), with a trend, though not significant, increase in lung weight normalized to tibia length in the NLC TAC mice (Fig. S2C). Together, these data demonstrate that βARKnt peptide expression preserves cardiac function during chronic pressure overload.

**Figure 2:**
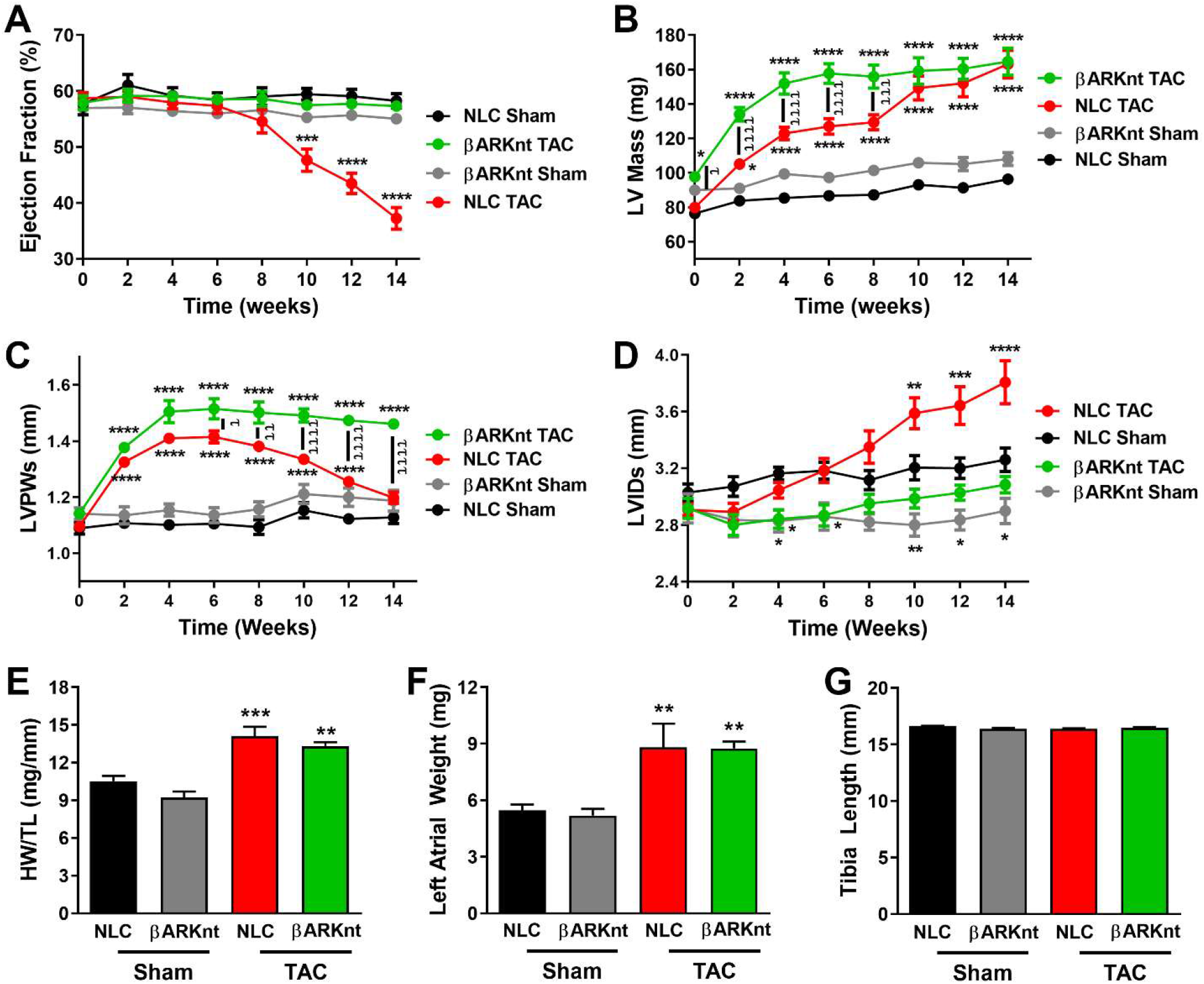
Despite enhanced hypertrophy, βARKnt hearts do not transition to heart failure after chronic pressure overload. Serial measures of non-transgenic littermate control (NLC) and TgβARKnt Sham and 14 week after TAC animals for (A) % left ventricular (LV) ejection fraction, (B) LV mass, (C) LV posterior wall thickness during systole (LVPWs), and (D) LV interior diameter during systole (LVIDs). *, p ≤ 0.05; **, p ≤ 0.01; ***, p ≤ 0.0007; ****, p < 0.0001 by two-way ANOVA with repeated measures and Tukey post-hoc test relative to corresponding NLC Sham. ^t^ p ≤ 0.03; ^tt^, p ≤ 0.006; ^ttt^, p = 0.0002; ^tttt^, p < 0.0001 by wo-way ANOVA with repeated measures and Tukey post-hoc test relative to corresponding NLC TAC. n = 9-15 mice per group. Measures of (E) heart weight normalized to tibia length (HW/TL), (F) left atrial weight, and (G) tibia length in these animals. **, p ≤ 0.01; ***, p = 0.0004 by one-way ANOVA with Tukey post-hoc test relative to NLC Sham. n = 9-15 mice per group.

To determine whether the enhanced hypertrophy and yet cardioprotection was consistent with inhibition of GRK2 activity at GPCRs we studied cardiac-targeted βARKct transgenic mice. The βARKct mice express the carboxyl-terminal 194 amino acids of GRK2 that competes for binding to G_βγ_ and membrane translocation, acting as a peptide inhibitor of GRK2 activity on GPCRs (Fig. 3A) [6]. Consistent with previous myocardial infarction studies, βARKct expression was cardioprotective, preventing the transition to heart failure 14 weeks post-TAC, as evidenced by the preserved cardiac ejection fraction, posterior wall thickness, and internal systolic diameter in the βARKct mice compared to control (Fig. 3B-D). Thus, the degree of cardioprotection observed in the βARKnt mice was equivalent with the peptide inhibition of GRK2 observed in the TgβARKct mice. We also studied mice with cardiac-restricted GRK2 overexpression, where the levels of GRK2 are similar to those observed in human heart failure (Fig. 3E) [13]. In contrast to the cardioprotection provided by βARKnt and βARKct, GRK2 overexpression demonstrated the opposite effect, hastening the decompensation to heart failure with a more rapid deterioration in function and a more progressive reduction in wall thickness and expansion of the LV (Fig. 3F-H).

**Figure 3:**
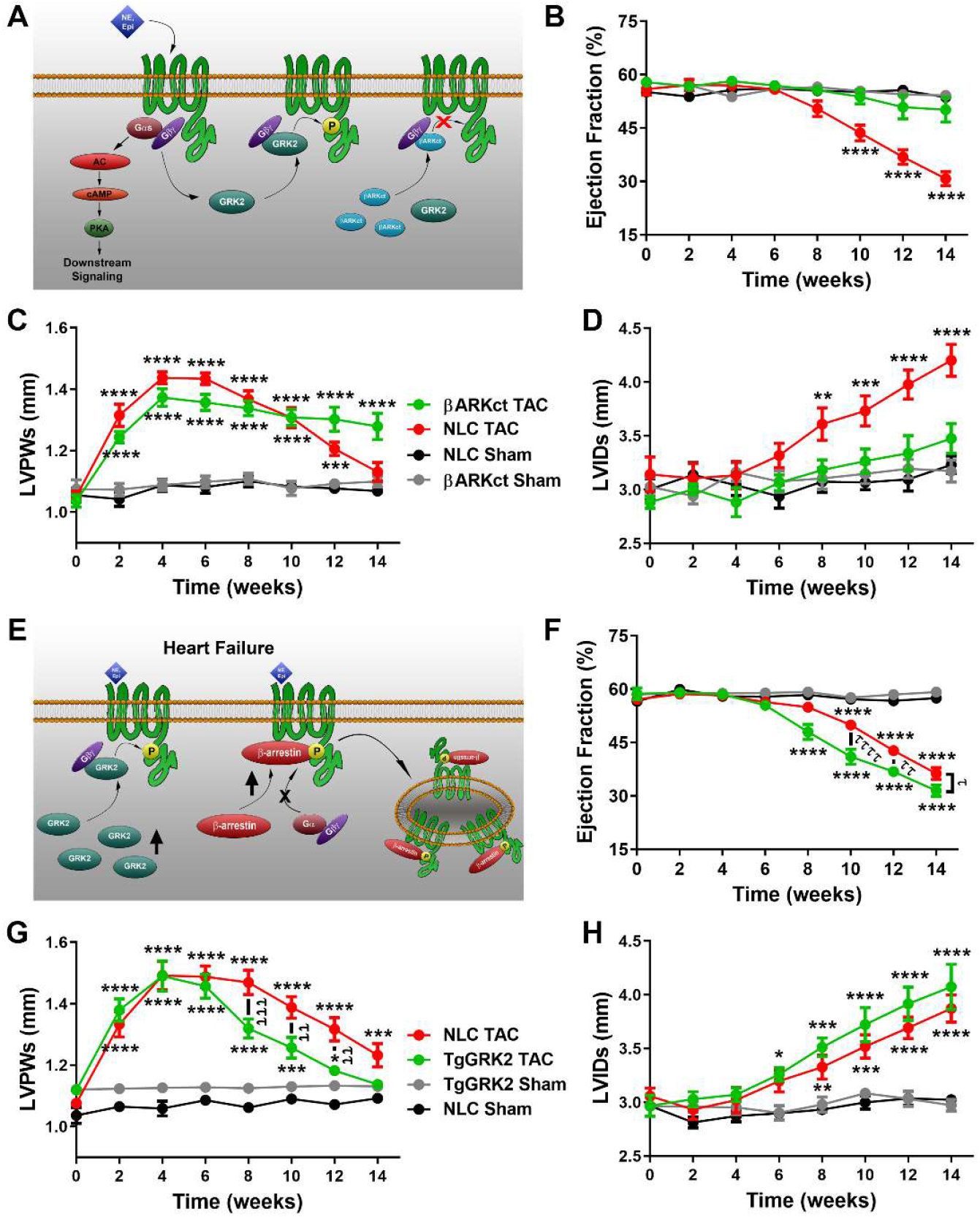
βARKnt elicits a similar degree of cardioprotection to that of βARKct, in contrast to the hastened transition to heart failure in the GRK2 overexpressing mice. (A) Cartoon representation of the intracellular activity of the βARKct peptide. Serial measures of non-transgenic littermate control (NLC) and TgβARKct Sham and 14 week after TAC animals for (B) % left ventricular (LV) ejection fraction, (C) LV posterior wall thickness during systole (LVPWs), and (D) LV interior diameter during systole (LVIDs). **, p = 0.0035; ***, p ≤ 0.0007; ****, p < 0.0001 by two-way ANOVA with repeated measures and Tukey post-hoc test relative to corresponding NLC Sham. n = 9-12 mice per group. (E) Cartoon representation of the intracellular effect of increased GRK2 expression. Serial measures of NLC and TgGRK2 Sham and 14 week after TAC animals for (F) % LV ejection fraction, (G) LVPWs, and (H) LVIDs. *, p ≤ 0.04; **, p = 0.0083; ***, p ≤ 0.0009; ****, p < 0.0001 by two-way ANOVA with repeated measures and Tukey post-hoc test relative to corresponding NLC Sham. ^t^ p ≤ 0.02; ^tt^, p ≤ 0.005; ^ttt^, p ≤ 0.0008; ^tttt^, p < 0.0001 by two-way ANOVA with repeated measures and Tukey post-hoc test relative to corresponding NLC TAC. n = 9-13 mice per group.

### 3.3. βARKnt Limits Adverse Ventricular Remodeling suggestive of Physiological Hypertrophy

To elucidate the underlying mechanisms by which βARKnt peptide expression enhances hypertrophy and yet elicits cardioprotection, tissues were taken for histological and biochemical analysis 4 weeks post-TAC. To visualize the effect of βARKnt peptide expression on myocardial structure, we performed wheat germ agglutinin staining to measure myocyte cross-sectional area in these mice. These data revealed a significant increase in myocyte size in transgenic Sham animals and an even greater, but equal, increase in NLC and βARKnt mice post-TAC (Fig. 4A, B), demonstrating an alteration in cardiomyocyte size prior to cardiac stress. Hypertrophic gene expression was measured via RT-PCR of cardiac mRNA and fetal gene induction was similar between these animals (Fig. 4C-E). These data demonstrate proportional hypertrophic growth in the TgβARKnt and non-transgenic mice in response to acute cardiac stress. To determine whether βARKnt expression was able to reduce maladaptive LV remodeling after TAC, fibrotic gene expression was measured via RT-PCR of cardiac mRNA. Matrix metalloproteinase-2, type III collagen, and TGFβ expression were all significantly reduced in TgβARKnt compared to non-transgenic mice, suggesting a decrease in cardiac fibrosis post-TAC (Fig. 4F-H). Masson Trichrome staining of four chamber paraffin sections taken at the level of the aortic outflow tract revealed a proportional increase in cardiac size in non-transgenic and TgβARKnt hearts 4 weeks post-TAC (Fig. 4J). To further investigate the integrity of the myocardium, we collected higher magnification images of the LV and observed a significant decrease in interstitial fibrosis in the βARKnt transgenic mice (Fig. 4I, K). Together, these data suggest that while βARKnt peptide expression does not appear to alter adaptive remodeling, it impedes maladaptive LV remodeling after acute pressure overload.

**Figure 4:**
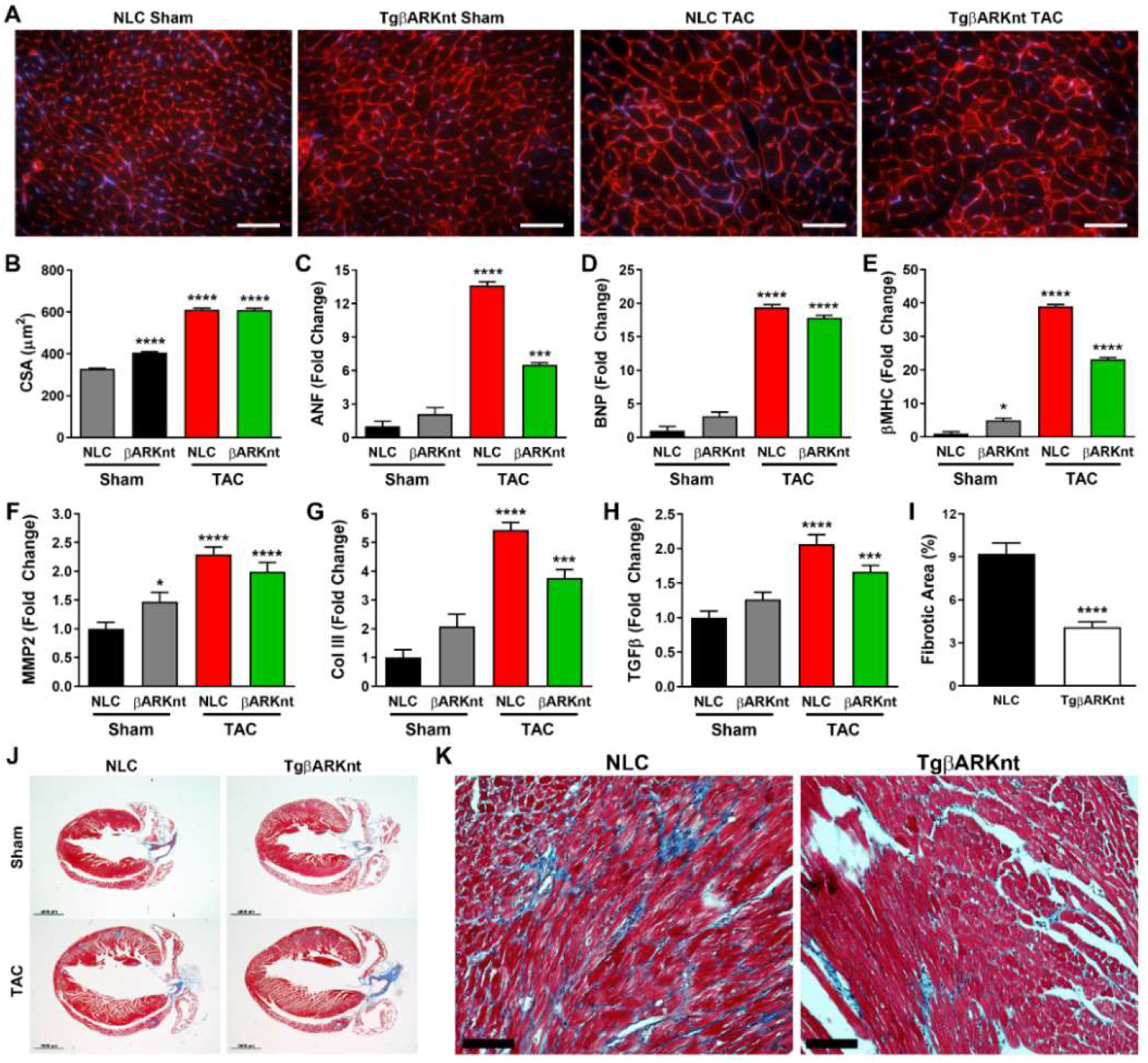
βARKnt mice exhibit preserved cardiomyocyte size, reduced neonatal and fibrotic gene expression, and reduced interstitial fibrosis 4 weeks after TAC. (A) Representative images of WGA and 4’,6-diamidino-2-phenylindole (DAPI)-stained murine heart sections from non-transgenic littermate control (NLC) and TgβARKnt hearts 4 weeks after sham or TAC surgery. Scale bar, 50 mm. (B) Quantification of cardiomyocyte cross-sectional area (CSA) in these animals. ****, p < 0.0001 by one-way ANOVA with Tukey post-hoc test relative to NLC sham. n = 5 to 10 hearts per group, 40 images per heart. Quantification of RT-PCR data showing fold change in (C) atrial natriuretic factor (*ANF*), (D) brain natriuretic peptide (*BNP*), and (E) β-myosin heavy chain *(βMHC),* (F) matrix metalloproteinase-2 *(MMP2),* (G) type III collagen (Col *III*), and (H) transforming growth factor beta (TGFβ) mRNA expression in these animals. *, p ≤ 0.018; ***, p ≤ 0.0006; ****, p < 0.0001 by one-way ANOVA with Tukey post-hoc test relative to NLC Sham. n = 11-12 mice per group. (I) Quantification of % fibrotic area from NLC and TgβARKnt hearts 4 weeks after TAC surgery. ****, p < 0.0001 by Student’s *t* test relative to NLC TAC mice. n = 191 images from 11 hearts for NLC mice (~16 images per heart) and 126 images from 9 hearts for TgβARKnt mice. (J) Representative images of Masson trichrome-stained murine heart sections in these mice. Scale bar, 2000 mm. (K) Representative higher magnification (×40) images of Masson trichrome-stained post-TAC murine heart sections from these mice demonstrating interstitial fibrosis. Scale bar, 200 mm.

To further understand the effect the βARKnt peptide on cardiac function we measured βAR density by radioligand binding and found that receptor density was preserved by βARKnt expression after 4 weeks of pressure overload (Fig. 5A). Further, there was a trend towards an increase in GRK2 levels in Sham animals, but this was restrained at 4 weeks (Fig. 5B). Given the significant thickening of the LV wall in the βARKnt mice at baseline and after cardiac stress combined with reduced fibrosis and cardioprotection during chronic stress we asked whether these hearts were more representative of a physiological hypertrophy. Interestingly, we found that despite a significant increase in total ERK in the βARKnt sham and 4 week post-TAC control hearts, normalized phosphorylated ERK was significantly increased only in the control TAC mice (Fig. S3A, Fig. 5C). Similarly, phosphorylated p38 levels were only elevated in control hearts 4 weeks after TAC, with no difference in total p38 levels (Fig. 5D, Fig. S3B). In contrast, with no difference observed in total levels across all groups, phosphorylated Akt levels were enhanced in the βARKnt TAC mice compared to control Sham and both the βARKnt Sham and TAC mice compared to control TAC (Fig. S3C, Fig. 5E). These data demonstrated that βAR regulation and cardiac signaling pathways are altered in the βARKnt transgenic hearts in a manner that suggests a beneficial signaling profile for cell survival and contractility that will require further investigation.

**Figure 5:**
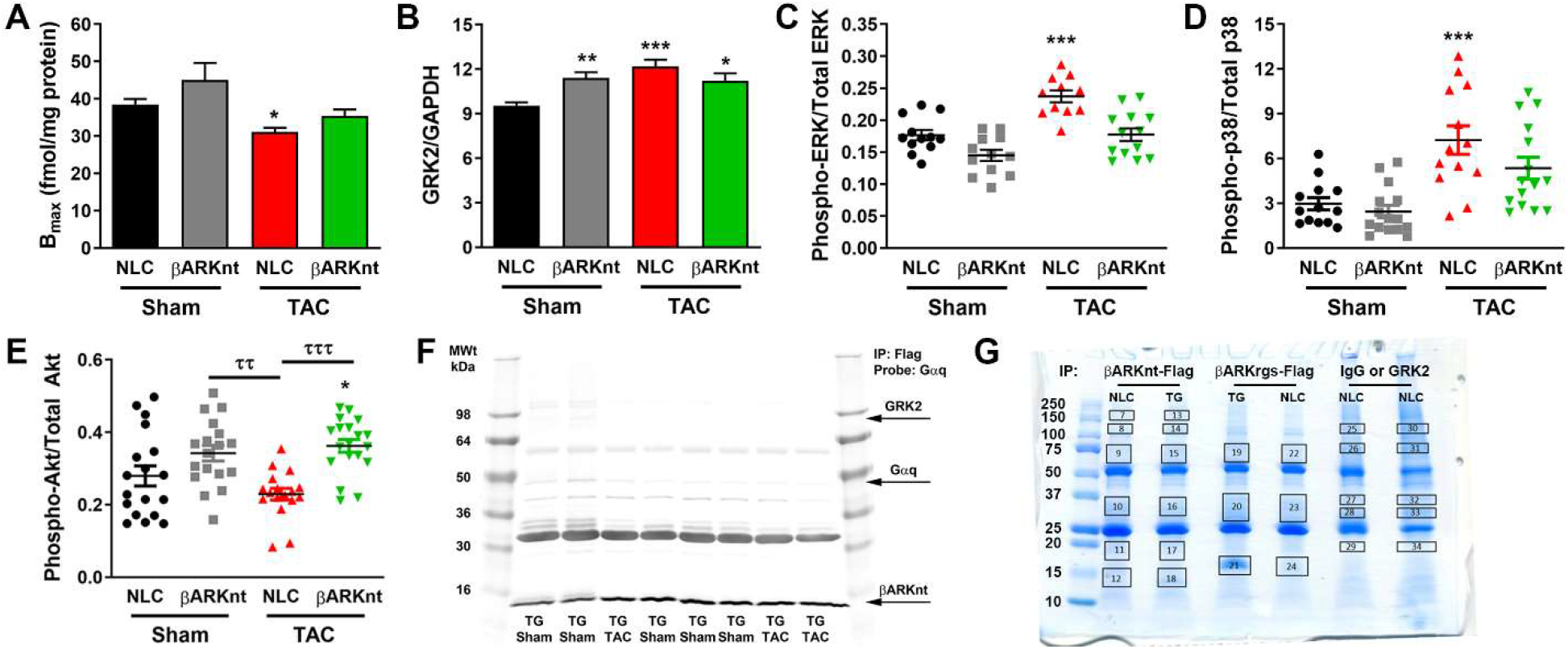
Interrogation of cardiac signaling pathways indicates a more physiological hypertrophy in the βARKnt mice and a distinct mechanism from the larger βARKrgs. (A) βAR density in non-transgenic littermate control (NLC) and TgβARKnt left ventricular tissue 4 weeks after Sham or TAC surgery (Bmax values shown as femtomoles of receptor per milligram of sarcolemmal protein). *, p = 0.043 by nonparametric one-way ANOVA with Dunn’s post-hoc test relative to NLC Sham. n = 7-12 mice per group. Quantification of (B) GRK2 normalized to GAPDH, (C) phosphorylated ERK normalized to total ERK, (D) phosphorylated p38 normalized to total p38, and (E) phosphorylated Akt normalized to total Akt in ventricular lysates from these animals. *, p ≤ 0.035; **, p = 0.0096; ***, p ≤ 0.0003 by one-way ANOVA with Tukey post-hoc test relative to NLC Sham. ^tt^, p = 0.0022; ^ttt^, p = 0.0002 by one-way ANOVA with repeated measures and Tukey post-hoc test relative to corresponding NLC TAC. n = 10-19 hearts each, from 5-9 Western blots. (F) Representative image of a monoclonal Flag immunoprecipitation (IP) followed by Western blotting probing for rabbit polyclonal GRK2, goat polyclonal Gαq, and rabbit polyclonal Flag-tagged βARKnt in cardiac lysates from NLC and TgβARKnt mice. (G) Coomassie-stained gel of IPs from cardiac lysates of TgβARKrgs-Flag or TgβARKnt-Flag mice or their corresponding NLCs using monoclonal Flag antibody-conjugated beads and IP of endogenous GRK2 from NLCs using a monoclonal GRK2 antibody-conjugated beads compared to mouse IgG-conjugated control beads, with boxes depicting areas that were cut from the gel lanes for extraction and sequencing by the Proteomics and Metabolomics Core.

### 3.4. βARKnt Acts Distinctly from βARKrgs and GRK2

To identify the protein interactions responsible for these beneficial cardiac effects induced by cardiac βARKnt expression, we performed immunoprecipitation (IP) reactions on cardiac lysates from NLC and TgβARKnt mice 4 weeks after Sham or TAC surgery. Unlike the previously published full βARKrgs peptide [1], IP of flag-tagged βARKnt did not co-immunoprecipitate (co-IP) with Gαq, demonstrating that they do not interact in complex with each other *in vivo* (Fig. 5F). Neither did βARKnt expression decrease IP3 activation after TAC (Fig. S3D). Further, βARKnt did not co-IP with full-length GRK2 *in vivo* (Fig. 5F, Fig. S3E). Western blots of cardiac lysates prior to and remaining after IP demonstrated that the flag IP was efficient and there was no difference in the amount of βARKnt IP’d from Sham or TAC animals (Fig. S3F, G). Together, these data demonstrate that the physiological effects of βARKnt are not due to altered Gαq-coupled GPCR activation or direct interference with GRK2 activity. We went on to interrogate the presumed GRK2 amino-terminal interacting partners and in contrast to *in vitro* data, were unable to confirm *in vivo* interactions with RKIP, clathrin, or Akt (Fig. S3H-J).

To overcome the limitations of conventional IP we performed a secondary method of proteomic analysis. We performed IPs on cardiac lysates from TgβARKrgs-Flag or TgβARKnt-Flag mice or their corresponding NLC mice using monoclonal Flag antibody-conjugated beads. In addition, we IP’d endogenous GRK2 from an NLC mouse using monoclonal GRK2 antibody-conjugated beads compared to mouse IgG-conjugated control beads. These samples were then run on a tris-glycine gel with each sample separated by an empty lane, and stained with Coomassie Blue (Fig. 5G). A total of 6 areas were cut from the NT Flag IP gel lanes, 3 from the RGS Flag IP, and 5 from the IgG and GRK2 IP gel lanes. Extraction and sequencing was performed by Dr. Belinda Willard, Director of the LRI Proteomics Laboratory, according to their standard procedures [14]. Briefly, these areas were digested with trypsin, the digests were analyzed with LC-MS/MS, and the data was searched against the mouse UniProtKB database. Over 600 proteins were identified in these samples. The protein of interest, GRK2, was identified as a major component of the Flag IP samples and the coverage maps for these proteins was consistent with the expression of N-terminal truncated forms. A comparative analysis of the Flag IP samples from control and βARKnt samples was performed along with the control and βARKrgs, and IgG and GRK2 IP samples, with a separate comparison for each area. Proteins demonstrated in the literature to bind nonspecifically to the most commonly used affinity matrices were excluded from analysis [15]. Proteins that were at least two-fold more abundant in the Flag/GRK2 IP samples that were also of moderate to high abundance (spectral count > 10) were highlighted as the best candidates for follow up experiments, and consisted of 15 hits for βARKnt (Table 1), 17 for βARKrgs (Table 2), and 31 for GRK2 (Table 3). Interestingly, the majority of hits were distinct for each target, supporting our hypothesis that the βARKnt peptide embodies distinct functional interactions that do not occur within the full-length enzyme *in vivo*, and elicit the enhanced hypertrophy and cardioprotection observed in these mice. Further, for a number of these hits the cardiovascular function has not been studied, perhaps due in part to a lack of appropriate tools for their study. For βARKnt, these hits included regulators of endocytic trafficking, metabolism, and mitochondria (Table 1). The significance of these interesting targets will be the subject of future studies.

**Table 2.** Summary for βARKrgs-Flag IP

|  | Protein | Accession | Gene | Area | Spectral Counts |  | SC Ratio |
| --- | --- | --- | --- | --- | --- | --- | --- |
|  |  |  | ID |  | Ln6 | Ln8 | Ln6/Ln8 |
| 1 | Atypical kinase COQ8A, mitochondrial | Q60936 | Coq8a | 1 | 15 | 0 | Ln6 only |
| 2 | Cytoskeleton-associated protein 4 | Q8BMK4 | Ckap4 | 1 | 12 | 0 | Ln6 only |
| 3 | T-complex protein 1 subunit theta | P42932 | Cct8 | 1 | 20 | 0 | Ln6 only |
| 4 | 3-ketoacyl-CoA thiolase, mitochondrial | Q8BWT1 | Acaa2 | 2 | 10 | 0 | Ln6 only |
| 5 | D-beta hydroxybutyrate dehydrogenase, mitochondrial | Q80XN0 | Bdh1 | 2 | 30 | 8 | 3.8 |
| 6 | Guanine nucleotide-binding protein G(I)/G(S)/G(T) subunit beta-2 | P62880 | Gnb2 | 2 | 10 | 0 | Ln6 only |
| 7 | Isoform 2 of Epimerase family protein SDR39U1 | Q5M8N4-2 | Sdr39u1 | 2 | 10 | 0 | Ln6 only |
| 8 | Mitochondrial coenzyme A transporter SLC25A42 | Q8R0Y8 | Slc25a42 | 2 | 16 | 0 | Ln6 only |
| 9 | Mitochondrial uncoupling protein 3 | P56501 | Ucp3 | 2 | 13 | 0 | Ln6 only |
| 10 | Prostaglandin E synthase 2 | Q8BWM0 | Ptges2 | 2 | 10 | 0 | Ln6 only |
| 11 | ADP-ribosylation factor 3 | P61205 | Arf3 | 3 | 12 | 0 | Ln6 only |
| 12 | Beta-adrenergic receptor kinase 1 | Q99MK8 | Grk2 | 3 | 395 | 7 | 56.4 |
| 13 | Calmodulin-3 | P0DP28 | Calm3 | 3 | 11 | 0 | Ln6 only |
| 14 | Cold-inducible RNA-binding protein | P60824 | Cirbp | 3 | 15 | 0 | Ln6 only |
| 15 | Microsomal glutathione S-transferase 3 | Q9CPU4 | Mgst3 | 3 | 12 | 0 | Ln6 only |
| 16 | NADH dehydrogenase [ubiquinone] 1 alpha subcomplex subunit 8 | Q9DCJ5 | Ndufa8 | 3 | 32 | 0 | Ln6 only |
| 17 | NADH dehydrogenase [ubiquinone] 1 beta subcomplex subunit 8, mitochondrial | Q9D6J5 | Ndufb8 | 3 | 15 | 0 | Ln6 only |
| 18 | Protein FAM162A | Q9D6U8 | Fam162a | 3 | 14 | 0 | Ln6 only |

**Table 3.**
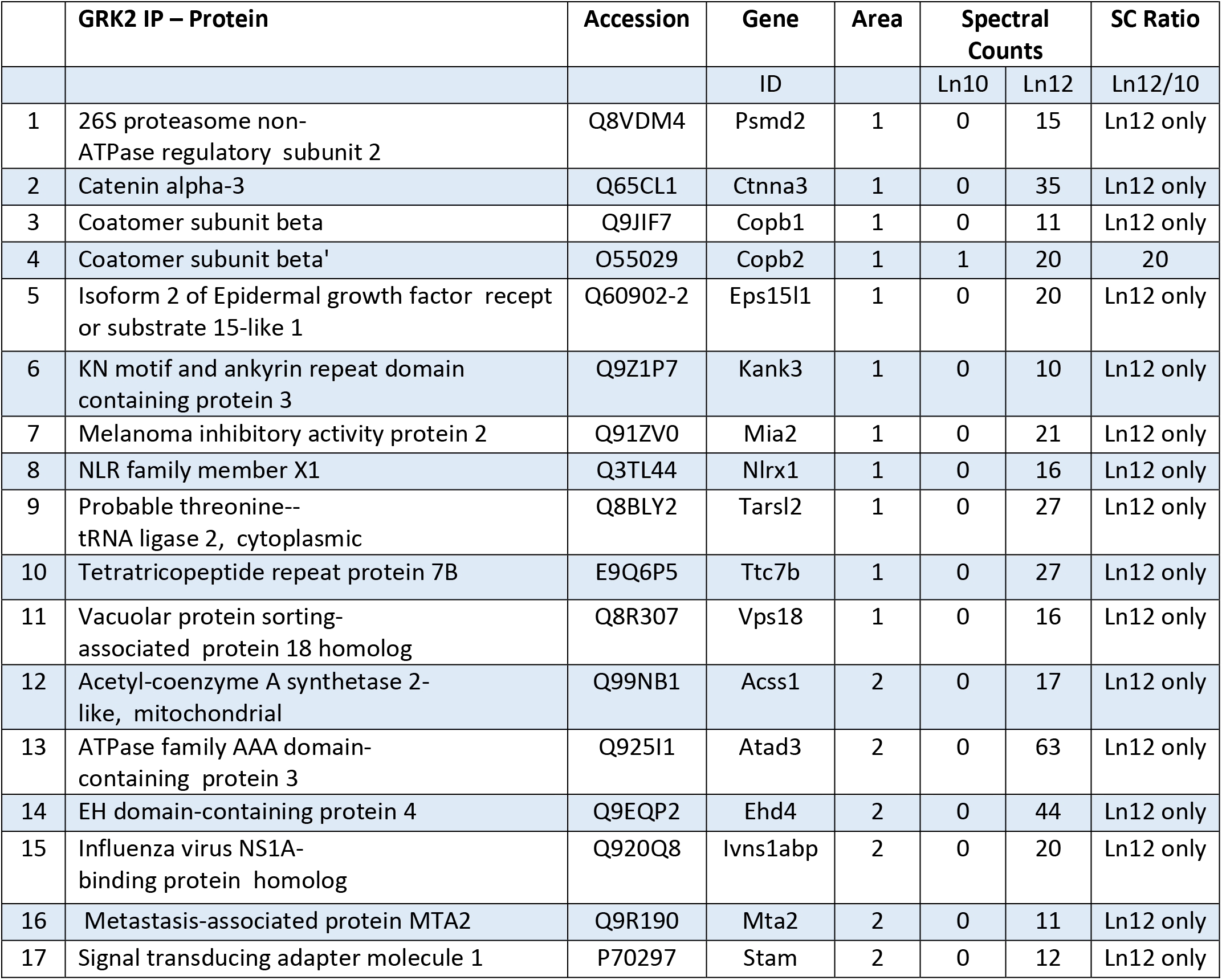
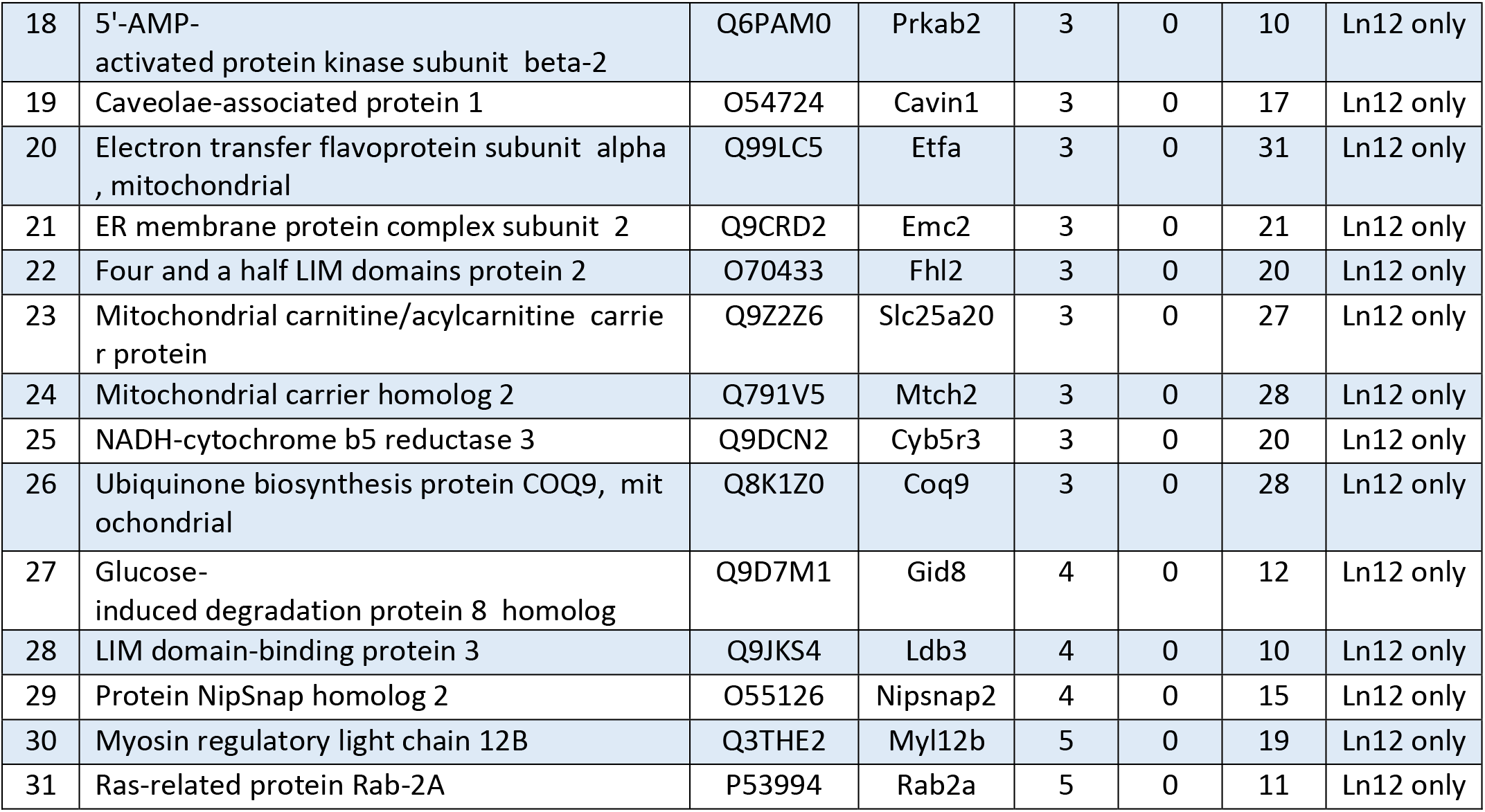
Summary for GRK2 IP

## 4. Discussion

Herein, we investigated whether the baseline hypertrophy and preserved βAR density exhibited in the βARKnt transgenic mice would alter cardiovascular remodeling and the transition to heart failure following pressure-overload injury. Cardiac-specific TgβARKnt mice exhibited normal hypertrophic growth following TAC, but maintained wall thickness and cardiac function during chronic pressure overload. TgβARKnt expression differed from overexpression within the context of the full GRK2 enzyme that has no effect on acute hypertrophy and yet hastens the transition to heart failure. Further, while the degree of cardioprotection is similar to that previously shown for βARKct, TgβARKnt expression still differs due to the baseline hypertrophy and proportional response to TAC leading to an enlarged, yet efficient heart. Further, βARKnt mice have less activation of fetal genes, reduced fibrosis, and improved survival signaling.

Together, these data suggest that the βARKnt mice may be undergoing physiological, rather than pathological hypertrophy. In both cases, the acute hypertrophic growth is an adaptive response to preserve cardiac function in response to injury or stress. Initially they share some common pathways, but as they progress, the underlying gene expression and biochemical changes enact vastly divergent effects on cardiac remodeling and functional outcomes, with preserved cardiac structure and function in physiological hypertrophy versus development of heart failure and/or arrhythmias and mortality in pathological hypertrophy. In fitting with the mild, 10-20% increase in cardiac mass and cardiomyocyte size that characterizes physiological hypertrophy, the βARKnt mice exhibit an approximately 12.6% increase in heart weight and 23.7% increase in myocyte cross-sectional area. Also consistent with physiological hypertrophy, contractile function is preserved and trending towards increased in these mice compared to control, both at baseline and following TAC, with no transition to heart failure and reduced interstitial fibrosis suggestive of improved myocyte survival. This is particularly important, as heart failure with preserved ejection fraction (HFpEF) is heart failure in the absence of a decrease in systolic function, but is often characterized by concentric hypertrophy, diastolic dysfunction, vascular rarefaction, and cardiac fibrosis [16–18]. Thus, the decrease in interstitial fibrosis in these animals is inconsistent with HFpEF or heart failure with reduced ejection fraction (HFrEF). Exercise tolerance is another method to distinguish between physiological hypertrophy and HFpEF, and will also be explored. Due to the chronic postnatal expression of βARKnt in these transgenic mice, it is not possible to investigate whether the baseline hypertrophy is reversible, a hallmark of physiological hypertrophy. This could be investigated in the future using a conditional model or adeno-associated viral expression of βARKnt.

As stated, the initial study of the βARKnt peptide showed that this fragment did not alter acute hypertrophy within 7 days after pressure overload or demonstrate RGS activity *in vivo* against Gq-mediated signaling. In contrast, βARKnt induced hypertrophy and elevated βAR density without altering agonist-induced contractility or adenylyl cyclase activity, due to a compensatory increase in GRK2 activity. Importantly, though, βAR downregulation in response to chronic agonist administration was attenuated by βARKnt expression. Herein, we showed that βAR downregulation in response to pressure overload was also attenuated by βARKnt. Together, these data indicate a novel regulation of βAR receptor density by βARKnt. Interestingly, activation of β1ARs has been shown to induce cardiac hypertrophy and atrial natriuretic factor transcription, and this hypertrophy is blocked by inhibition of each step in the signaling cascade, including β1AR, βARK1 (GRK2), β-arrestin1, Src, dynamin, and the endocytic machinery [19, 20]. These studies demonstrated that activation of the endocytic machinery plays a particularly important role. Data from the initial study showing that the nonselective βAR antagonist nadolol reversed the baseline hypertrophy and reduced membrane translocation of GRK2, suggests the importance of chronic βAR signaling in the βARKnt mice to the phenotypical characterization. Though βARKnt expression does not alter adenylyl cyclase activity downstream of βAR activation, nor does it enhance atrial natriuretic peptide expression, numerous regulators of endocytic trafficking were identified in our proteomics search. These data suggest that he βARKnt-mediated regulation of βAR density may provide a novel means of cardioprotection during pressure-overload induced cardiac dysfunction and failure.

Interestingly, the reactivation of fetal genes including ANP, BNP, and βMHC was mildly enhanced in the βARKnt Sham animals and only ANP and βMHC, not BNP, were slightly reduced in these mice after TAC. Induction of these fetal genes is characteristic in pathological hypertrophy, whereas their expression is generally unaltered or decreased during physiological hypertrophy. Coordinate with gene induction are changes in biochemical signaling within the cardiomyocyte, and the culmination of these signaling pathways determines physiological versus pathological hypertrophy [21–25]. Data regarding the role of ERK1/2 signaling in hypertrophic cardiac remodeling is complex, showing both beneficial and detrimental effects that are likely dependent upon spatiotemporal activation and upstream signaling, but there is a consensus that ERK signaling contributes to cardiac hypertrophy through regulation of nuclear transcription factors [26]. Additionally, the GATA4 transcription factor that promotes pathological activity is activated in response to phosphorylation by p38 kinases and JNKs [26]. Of these mitogen-activated protein kinase pathways, both ERK1/2 and p38 were significantly enhanced in control mice 4 weeks after TAC and this elevation was abrogated by βARKnt expression. Coordinately, cell survival signaling is critical for ensuring physiological, not pathological hypertrophy. In contrast to ERK1/2 and p38, phosphorylated Akt was enhanced in the βARKnt Sham and TAC mice compared to post-TAC NLC mice. Interestingly, phosphorylated/activated Akt is not only a marker of enhanced survival signaling, but a possible indicator of improved insulin signaling in the heart.

In cardiomyocytes, insulin binding to the insulin receptor (InsR) signals through insulin receptor substrate (IRS) and phosphoinositide 3-kinase (PI3K) to generate phosphatidylinositol 3,4,5-trisphosphate and activation of the phosphatidylinositol-dependent kinases PDK1 and PDK2. Both of these activities lead to the serine/threonine phosphorylation and activation of protein kinase B (PKB, i.e. Akt), which then targets Akt substrate of 160 kilodaltons (AS160) [27, 28]. AS160 is a Rab GTPase-activating protein (GAP), and it’s phosphorylation by Akt inhibits this GAP activity, allowing an increase in the GTP-bound active Rab and translocation of Glut4 containing vesicles to the plasma membrane [29–32]. Akt phosphorylation also inhibits glycogen synthase kinase 3β (GSK3β), thereby releasing its inhibition of the translation initiation factor eIF2Bε and allowing for beneficial transcriptional activation. Interestingly, we observed a significant increase in phosphorylated GSK3β in the βARKnt Sham mice (data not shown), suggesting that this pathway may be active at baseline in these mice and provided a more amicable substrate for adaptive versus maladaptive remodeling. Physiological hypertrophy is marked not only by an increase in cell survival signaling, but also increased and efficient energy production, antioxidant regulation, and enhanced mitochondrial quality control that antagonize pathological remodeling to produce a purely adaptive response. Further, not only changes in cardiomyocyte, but systemic metabolism, precede heart failure development [33–37]. This is particularly relevant since AS160 (Tbc1d4) was identified as one of the hits for βARKnt. Thus, an inquiry into the effect of βARKnt on cardiomyocyte and systemic metabolism will be a major focus of future studies into the potential protective mechanism of this protein fragment.

In this study we immunoprecipitated βARKnt, βARKrgs, and GRK2 and performed proteomic analysis. This study focused on bands of interest where differential protein expression was observed by Coomassie staining, and identified numerous distinct hits for each target protein. As stated above for βARKnt, these hits included regulators of endocytic trafficking, metabolism, and mitochondria. It is known that GRK2 translocates to and disrupts mitochondrial function during cardiac stress by reducing fatty acid oxidation efficiency [38–40]. However, the subcellular localization of βARKnt within the myocyte at baseline or following cardiac stress is, as of yet, unknown. Confirmation of these mitochondrial hits and investigating whether βARKnt alters mitochondrial function will be part of our future metabolic studies.

In summary, we found that transgenic expression of the short N-terminal domain of GRK2, βARKnt, demonstrates a pro-hypertrophic effect at baseline, yet proportional hypertrophic growth and cardioprotection in a murine model of chronic pressure overload. This increased wall thickness and absence of heart failure development was not replicated by βARKct or GRK2 overexpression. Further, βARKnt preserved LV structure, reduced interstitial fibrosis, and enhanced cell survival signaling in the heart in a way that, despite the as yet unknown mechanism, suggests physiological hypertrophy. While ongoing studies are needed to uncover the mechanism of action of βARKnt, the identification of regulators of protein trafficking and metabolism in our proteomic study suggest key future directions for this project. βARKnt-mediated regulation of βAR density may provide a novel means of cardioprotection during pressure-overload induced heart failure. Further, this line may represent a unique tool to interrogate the underlying mechanisms of physiological versus pathological hypertrophy, and thus a means to identify novel therapeutic modalities for the treatment of human heart failure.

## Supporting information

Schumacher 2020 Supplement

## Abbreviations

GPCR: G protein-coupled receptor;
GRK2: G protein-coupled receptor kinase 2;
RGS: regulator of G protein Signaling domain;
βAR: beta-adrenergic receptor;
LV: left ventricular;
TAC: transverse aortic constriction;
αMHC: α-myosin heavy chain;
NLC: non-transgenic littermate control;
WGA: wheat germ agglutinin;
IP: immunoprecipitation;
IP3: Inositol 1,4,5-trisphosphate;
^125^I-CYP: [^125^I]cyanopindolol;
HFpEF: heart failure with preserved ejection fraction;
HFrEF: heart failure with reduced ejection fraction;
InsR: insulin receptor;
IRS: insulin receptor substrate;
PI3K: phosphoinositide 3-kinase;
PDK1 and 2: phosphatidylinositol-dependent kinases;
PKB/Akt: protein kinase B;
AS160: Akt substrate of 160 kilodaltons;
GAP: GTPase-activating protein;
GSK3β: glycogen synthase kinase 3β.

## Acknowledgements

We thank Dr. Belinda Willard, Director of the LRI Proteomics Laboratory, for gel extraction, LC-MS/MS, sequencing and analysis of the βARKnt, βARKrgs, and GRK2 interacting partners.

