## Supplementary material for "A Peptide of the Amino-Terminus of GRK2 Induces Hypertrophy and Yet Elicits Cardioprotection after Pressure Overload": Schumacher 2020 Supplement: Schumacher 2020 Supplement.docx

**Supplemental Materials**

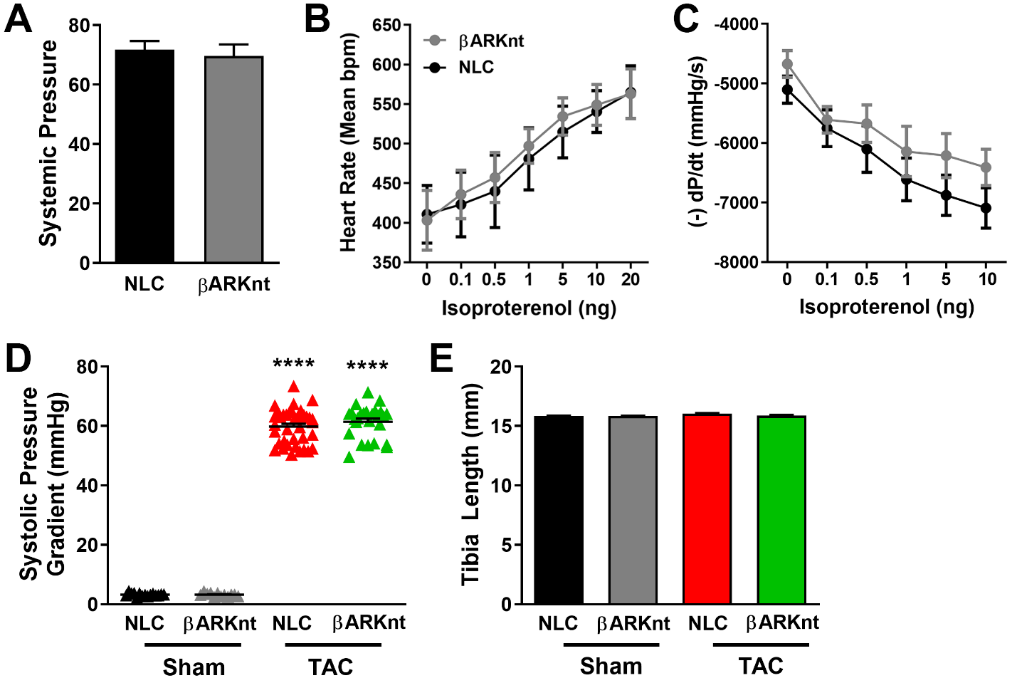

**Figure S1:** *Cardiac-specific βARKnt expression elicits baseline hypertrophy and additive pressure overload-induced hypertrophy.* Hemodynamics were recorded from 10-12 week old TgβARKnt and non-transgenic littermate control (NLC) mice. Quantification of (A) mean systemic pressure, (B) heart rate, and (C) LV =dP/dt average minimum at baseline and with increasing doses of isoproterenol (0.1-10 ng). n = 12 mice per group. (D) Measures of the systolic pressure gradient in NLC and TgβARKnt Sham and post-TAC mice 1 week after surgery. (E) Measures of tibia length in these animals 4 weeks after surgery. ****, p < 0.0001 by one-way ANOVA with Tukey post hoc test relative to NLC Sham. n = 37-62 mice per group.

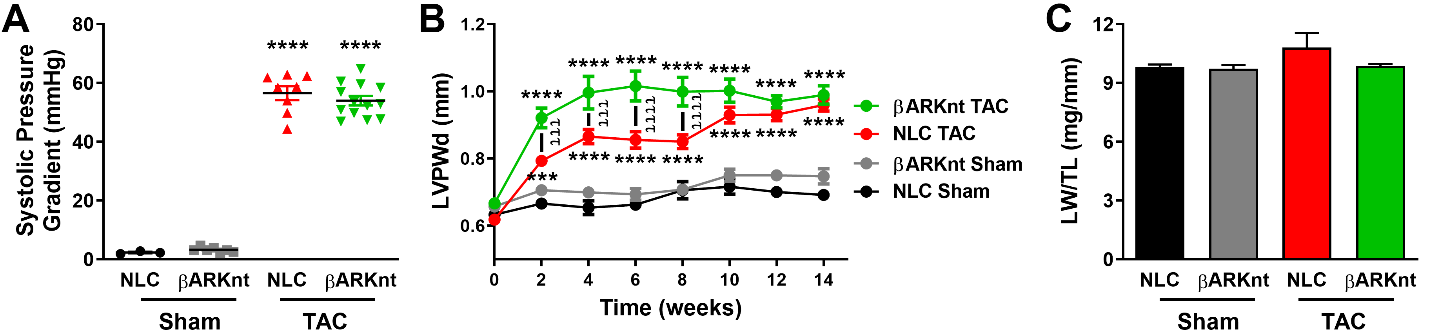

**Figure S2:** *Despite enhanced hypertrophy, βARKnt hearts do not transition to heart failure after chronic pressure overload.* (A) Measures of the systolic pressure gradient in non-transgenic littermate control (NLC) and TgβARKnt Sham and post-TAC mice 1 week after surgery. (B) Serial measures of left ventricular (LV) posterior wall thickness during diastole (LVPWd) in these mice. ***, p = 0.0008; ****, p < 0.0001 by one-way ANOVA with Tukey post hoc test relative to NLC Sham. ^ttt^, p ≤ 0.0005; ^tttt^, p < 0.0001 by one-way ANOVA with repeated measures and Tukey post-hoc test relative to corresponding NLC TAC. (C) Measures of lung weight normalized to tibia length (LW/TL) in these animals 14 weeks after surgery. n = 9-15 mice per group.

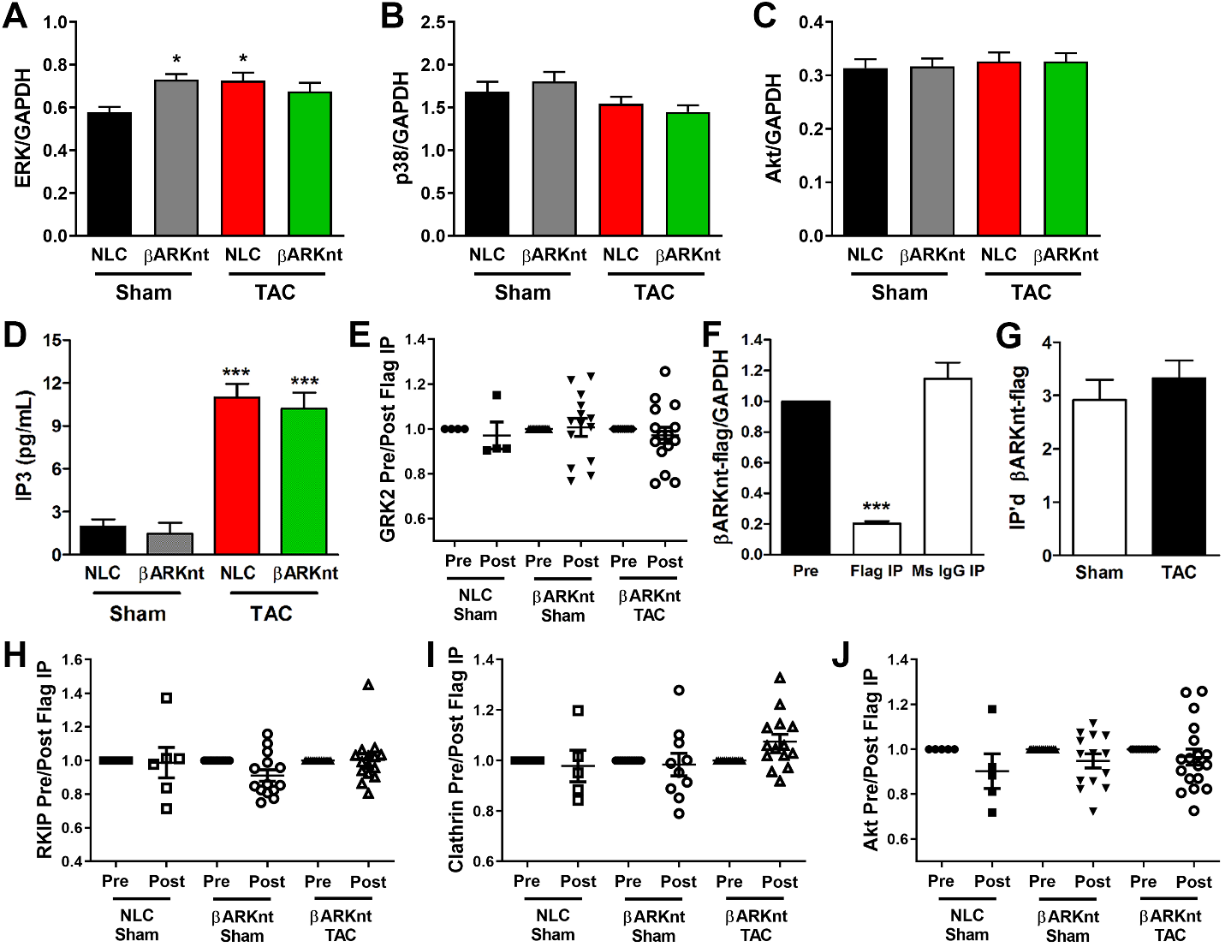

**Figure S3:** *Interrogation of cardiac signaling pathways indicates a more physiological hypertrophy in the βARKnt mice and a distinct mechanism from the larger βARKrgs.* Quantification of (A) total ERK normalized to GAPDH, (B) total p38 normalized to GAPDH, and (C) total Akt normalized to GAPDH in non-transgenic littermate control (NLC) and TgβARKnt lysates 4 weeks after sham or TAC surgery. *, p ≤ 0.017 by one-way ANOVA with Tukey post-hoc test relative to NLC Sham. n = 10-19 hearts each, from 5-9 Western blots. (D) Quantification of IP3 concentrations from NLC and TgβARKnt lysates 4 weeks after sham or TAC surgery. ****, p < 0.0001 by one-way ANOVA with Tukey post-hoc test relative to NLC Sham. n = 6-14 mice per group. (E) Quantification of GRK2 from Western blot samples collected pre and post IP of Flag in NLC and TgβARKnt lysates 4 weeks after Sham or TAC surgery, normalized first to GAPDH and then to pre-IP density. n = 4-15 hearts each, from 3-5 Western blots with Sham and TAC hearts run side by side. (F) Quantification βARKnt-Flag from Western blot samples collected pre and post IP for mouse monoclonal Flag or mouse IgG control in TgβARKnt lysates 4 weeks after Sham or TAC surgery, normalized first to GAPDH and then to pre-IP density. ****, p < 0.0001 by one-way ANOVA with Tukey post-hoc test relative to pre-IP control. n = 5-20 hearts each, from 5-7 Western blots with Sham and TAC hearts run side by side. (G) Quantification of total βARKnt-Flag in TgβARKnt lysates after monoclonal Flag IP, 4 weeks after sham or TAC surgery. n = 11-12 hearts each, from 3-5 Western blots. Quantification of (H) RKIP, (I) clathrin, and (J) Akt from Western blot samples collected pre and post IP of Flag in NLC and TgβARKnt lysates 4 weeks after Sham or TAC surgery, normalized first to GAPDH and then to pre-IP density. n = 5-18 hearts each, from 3-6 Western blots with Sham and TAC hearts run side by side.
